# Binary integer programming for Multiple Sequence Alignment

**DOI:** 10.1101/854786

**Authors:** S. Ali Lajevardy, Mehrdad Kargari

## Abstract

Molecular biology advances in the past few decades have contributed to the rapid increase in genome sequencing of various organisms; sequence alignment is usually considered as the first step in understanding the molecular function of a sequence. An optimal alignment adjusts two or more sequences in a way that it could compare the maximum number of identical or similar residues. The two sequence alignments types are: Pairwise Sequence Alignment (PSA) and Multiple Sequence Alignment (MSA). While dynamic programming (DP) technique is used in PSA to provide the optimal method, it will lead to more complexity if used in MSA. So, the MSA mainly uses heuristic and approximation methods. This paper presents a mathematical model for MSA that can be used as a basis for optimal solution in different ways. In order to obtain the results, the model is implemented using Genetic Algorithm method on the web.

## 1. Introduction

Bioinformatics uses biology, computer science, mathematics and statistics to analyze and interpret biological data [1]. In the last few decades, advances in molecular biology and development of the equipment in this field have led to a rapid increase in the genome sequencing of many species. This has extended to a point that genome sequencing projects have now become very common, and sequencing (SA) is usually considered as the first step in understanding the molecular phylogeny of an unknown sequence in bioinformatics [2]. This is done by aligning an unknown sequence with one or more known sequences in the database to predict the parts of sequences that could lead to performing evolutionary functional and structural roles [3]. An optimal alignment adjusts two or more sequences in order to compare the maximum number of identical or similar residues [4]. Sequences can be of two types: nucleotide sequences (DNA or RNA) or amino acid sequences (proteins).

The rearrangement process may create one or more spaces or gaps in alignment. A space indicates the probable loss or increase of a residue, thus the degree of evolution or deletion (in-del), as well as the transfer and reversal in SA is observed. Two types of sequence alignment are: Pairwise Sequence Alignment (PSA) and Multiple Sequence Alignment (MSA). The PSA aligns two sequences, while the MSA aligns several (more than two) related sequences. The benefits of MSA are greater than those of PSA, because MSA considers several members of the same family of sequences, therefore it provides more biological information. Besides, MSA is also a prerequisite for comparative genome analysis to identify and measure the conserved parts or functional points in a complete sequence family and to estimate evolutionary deviation between the sequences and even for ancestral sequence profiles [5]. Sequencing at the amino acid level is more related to the nucleotide level because the protein is the key to the functionality of biological molecules, hence it carries structural or functional information [6]. So, alignment is strongly linked to structural biology [7]. Therefore, sequence alignment, especially MSA, is the starting point for any field of biological research. MSA is deemed as a window for viewing the evolutionary, functional, and structural perspective of biological macromolecules in a concise format [8].

Different methods are used for scoring the sequence alignment to identify their similarity. Nucleotide scoring matrix is a simple identification scheme in which identical positions in both sequences yield positive scores. On the other hand, for a protein, a similarity score matrix is also counted (along with the identical score), which represents an amino acid with similar physical and chemical properties.

Replacement matrices that are often used for computing protein sequences include Point Accepted Mutation (PAM) [9] and BLOSUM [10]. The SA techniques can be of two types: Global Alignment and Local Alignment [12]. Global alignment is applied when the similarity is counted throughout the whole sequence. Several MSA techniques perform global alignment [13, 14]. However, problems arise when sequences are matched only in local regions, where there is a clear block of shared discontinuities across all sequences or altered domains exist among related sequences. In such cases, Local Alignment technique is adopted to identify similar regions among sequences [15, 16, 17]. When there is a great deal of variation along the sequences to be compared, local coordination is usually performed [18]. Figure (1) shows the difference between the two alignments.

**Fig 1.**
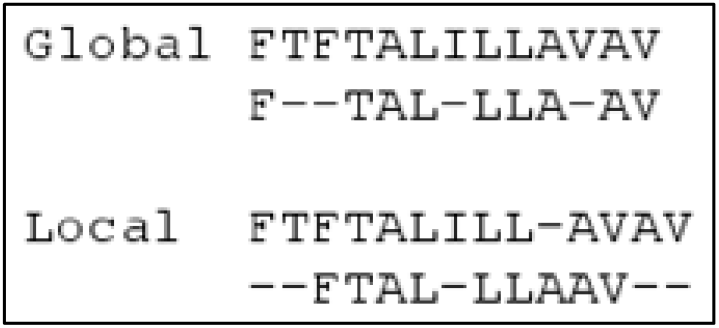
Difference between Global and Local Alignment.

For PSA, dynamic programming (DP) always provides the optimal orientation for a given objective function by finding the optimal way of alignment. It does this through the tracking process [19, 20]. The objective function is used to evaluate the alignment quality of a set of input sequences.

A lot of computational problems occur in these techniques, which require many computational resources [21]. In addition, dynamic programming faces major problems in MSA because the number of sequences is equal to the number of dimensions. If two or more optimal paths are available and require a tracking process, the complexity increases and the calculation of an accurate MSA becomes NP-Complete; therefore, the exact method in has been used in MSA for unrealistic small datasets [21].

The usual MSA methods are therefore heuristic or approximate, which provide a possible solution over a short and limited period of time. The existing heuristics do not provide the best solution. Besides, due to the high speed of database size growth, the development of new techniques for MSA is still under investigation [2].

## 2. Multiple Sequence Alignment Methods

Due to the complex relationship between related sequences and sometimes because of the lack of evolutionary history, most MSAs can hardly determine the final sequence [18]. Three types of alignment are often used in MSA: accurate, progressive, and iterative. Accurate algorithms usually deliver a high quality result that is very close to the desired endpoint [22, 23]. These methods attempt to arrange multiple sequences simultaneously and thus require dynamic programming. Other methods include tree alignment [24, 25] and star alignment [26, 27] and progressive alignment [28, 29]; however, these methods will not be effective when the sequence length increases [30].

### 2.1. Progressive Alignment

The most commonly used method applies a heuristic technique (hierarchical or tree method) for aligning multiple sequences. Such technique produces the final MSA by combining the two-pair alignment that starts with the most similar pair and continues to the farthest pair. All progressive alignments require two steps: the first step in which the relationship between the sequences is represented by a tree called the guide tree, and the second step in which the MSA is given based on the guide tree and linking sequences to each other. Figure (2) is an example of tree alignment [2].

**Fig 2.**
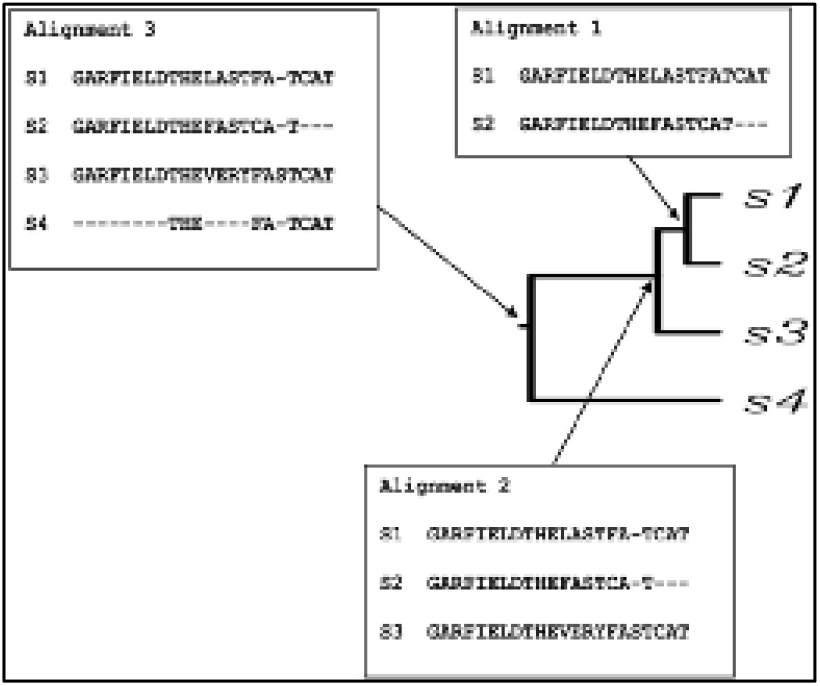
Tree alignment.

The primary guide tree is determined by an efficient clustering method such as neighbor-joining or UPGMA [31]. Progressive alignment cannot be optimal overall. The main problem is that when errors occur in any of the build steps of MSA, they find their way through the final stage. The performance also degrades significantly when all of the sequences in the set are rather distantly related. A very popular progressive alignment method is the Clustal family [32], especially the weighted variant ClustalW. The ClustalW method is widely used for phylogenetic tree construction. Another common progressive alignment method is called T-Coffee [33], which is slower than Clustal and its derivatives, but generally produces more accurate alignments for distantly related sequence sets.

### 2.2. Iterative Alignment

A set of methods to produce MSAs which reduce the errors inherent in progressive methods are classified as iterative because they work similarly to progressive methods but repeatedly realign the initial sequences as well as adding new sequences to the growing MSA [34]. One reason progressive methods are so strongly dependent on a high-quality initial alignment is the fact that these alignments are always incorporated into the final result - that is, once a sequence has been aligned into the MSA, its alignment is not considered further. This approximation improves the efficiency at the cost of accuracy. By contrast, iterative methods can return to previously calculated pairwise alignments or sub-MSAs containing subsets of the query sequence as a means of optimizing a general objective function such as finding a high-quality alignment score [31]. A well-known alignment method based iteration is the MUSCLE alignment that uses a more accurate distance measurement criterion to calculate the degree of sequence relevance [35]. The distance updated per round.

### 2.3. Hidden Markov Model

Hidden Markov Models (HMMs) are probabilistic models that can assign likelihoods to all possible combinations of jumps, matches, and mismatches to determine the most likely MSA or set of MSAs. HMMs can produce only one scoring at the highest level, but they can also produce a family of biologically significant alignments. HMMs can produce local and global alignments. Although HMM-based methods have are relatively new, they have made remarkable improvements in computational speed, especially for the sequences that contain overlapping regions [31]. Typically, HMM-based methods work by displaying the MSA as a directional non-circular graph containing a series of nodes representing possible columns for an MSA. The display of a definitely conserved column (meaning that all the sequences of an MSA share the same character in a particular location) is coded as a node that relates the number of possible characters from the next alignment column [36]. Numerous software applications have been implemented for a variety of HMM-based methods and are considered scalable and efficient, although the correct use of HMM is far more complex than conventional progressive methods. The simplest of these programs is Partial-Order Alignment (POA); a similar but more general approach has been implemented in SAM (Sequence Alignment and Modeling System) [37] and HMMER [38] packages.

### 2.4. Genetic Algorithm and Simulated Annealing Algorithm

Genetic algorithms have been used for MSA production in an attempt to simulate an evolutionary process that gave rise to the divers and divergent data in the query set. The method works by breaking a series of possible MSAs into fragments and repeatedly rearranging those fragments with the introduction of gaps at varying positions. A general objective function is optimized during the simulation. Generally, it is the “sum of pairs” function and is aimed to be maximized.

A technique has been implemented for protein sequences in the software program ‘SAGA’ (Sequence Alignment by Genetic Algorithm) and its equivalent for RNA has been implemented in RAGA. The technique of simulated annealing starts its work with an existing MSA which has been produced by another technique, and uses a series of rearrangements designed to find more optimal regions of alignment space than the one the input alignment already occupies. The simulated annealing algorithm, just like the genetic algorithm, maximizes an objective function (e.g. the sum-of-pairs function). Simulated annealing has been implemented in the program MSASA (Multiple Sequence Alignment by Simulated Annealing).

## 3. Mathematical Modeling of MSA

As mentioned in the review of the MSA methods, no mathematical model was found to optimize the MSA, and in this study, we are trying to present a mathematical model for multiple sequence alignment.

Sequence alignment is usually defined as an optimization problem where the goal is to find the best match with the highest score (minimum negative score). The way an alignment is scored is done based on the number and types of the residues that match each other.

The set of sequences chosen for alignment can be expressed through the following relation (1).

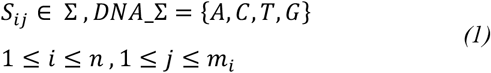

In this regard, S_ij_ represents the residues of the sequence i at position j, where the length of the strings is equal to i times of m_i_ and the number of strings is n (n represents the inputs to the model). These residues are members of the set Σ. Four of these residues, namely A, C, T, G (Adenine, Cytosine, Thymine and Guanine) exist for the DNA. Next, we create the string S′ on the basis of every binary x_ik_ element. This string is the conversion of the input S string into a string with different gaps based on the matrix X, which is defined in the following relation (2):

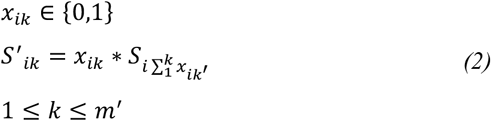

In this relation, S′_i_ is a sequence constructed on the basis of the Si sequence, which itself is constructed on the basis of the matrix X. The X is a zero-one matrix.

The construction the new string S′ _i_ is done from S_i_ with k length. The k length contains the “−” character as a gap. The “*” is created in a way that if x_ik_ is equal to zero, the “−” is built, otherwise the same character S_ij_ will be substituted.

The important point in this relation is that all the possible sequences can be created by matrix X; however, the m′ is determined based on other limits.

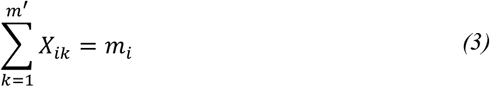

In Relation (3), the main limit is explained, as the number of 1s in each row should be equal to the length of m_i_.

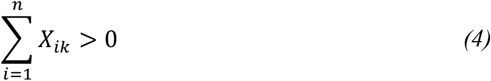

In this relation, regarding the need to be at least one final sequence at the designated position other than the element gap, the sum of 1s in each column of the matrix X must be bigger than zero.

In this case, m′ will be determined by the relations (5) and (6).

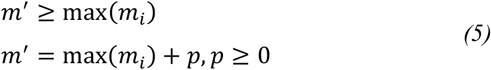

m′ must be equal to or bigger than the length of the input sequences. In other words, m′ must be p bigger than the maximum length of the sequence. p value is bigger than or equal to zero.

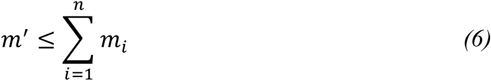

Given the relation (4), m′ must be equal to or less than the sum of the lengths of the input sequences. In case the length of m′ is equal to the sum of the lengths of the input sequences, the final solution is definitely the optimal one.

Finally, based on the defined variables and limitations, the minimizing function will be in the form of relation (7).

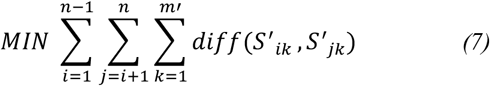

In the above-mentioned relation, the diff can of any type of the functions that were explained for scoring in the previous parts.

## 4. Implementation and Results

This model has been implemented by PHP language. It is available at: http://oms.modares.ac.ir/Genome/MSA/.

As shown in Fig. 3, the input sequences can be of any length and number. Besides, the model can be implemented based on the mismatch penalty and gap penalty as well as the parameter p - according to Relation 5, the P value determines the value of m′. The Genetic algorithm has been used for solving the problem, which gives optimal answers according to the population, number of generations, mutation coefficient and intersection factor.

**Fig 3.**
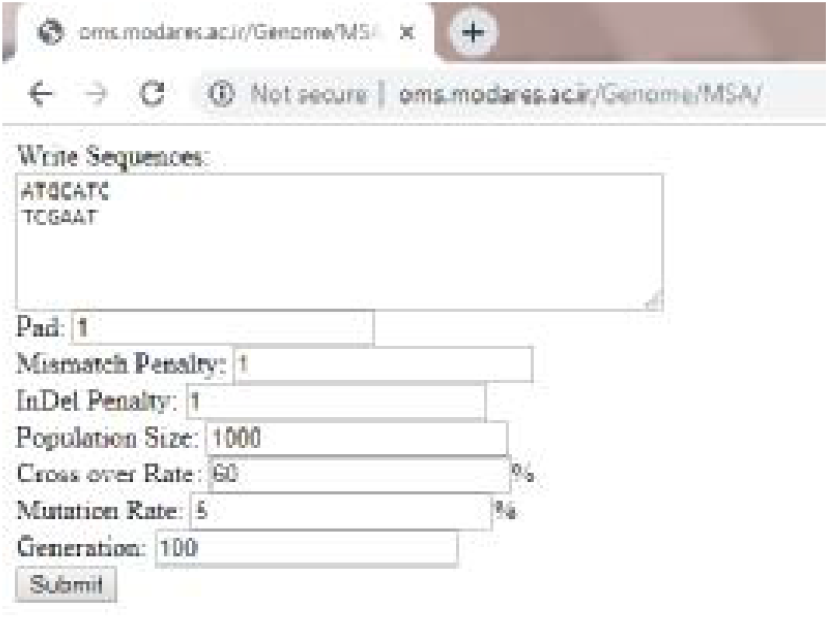
Implement MSA model on web.

It should be noted that in this method, the optimal sequences obtained are both global and local alignments.

Some results have been presented in Table (1). In each row, the three final sequence sets are based on sequence input, p value = 3, the gap penalty and substitution = 1, with a population of 1,000 and generation of 100. The cost of each obtained set is shown in C.

**Table 1.**
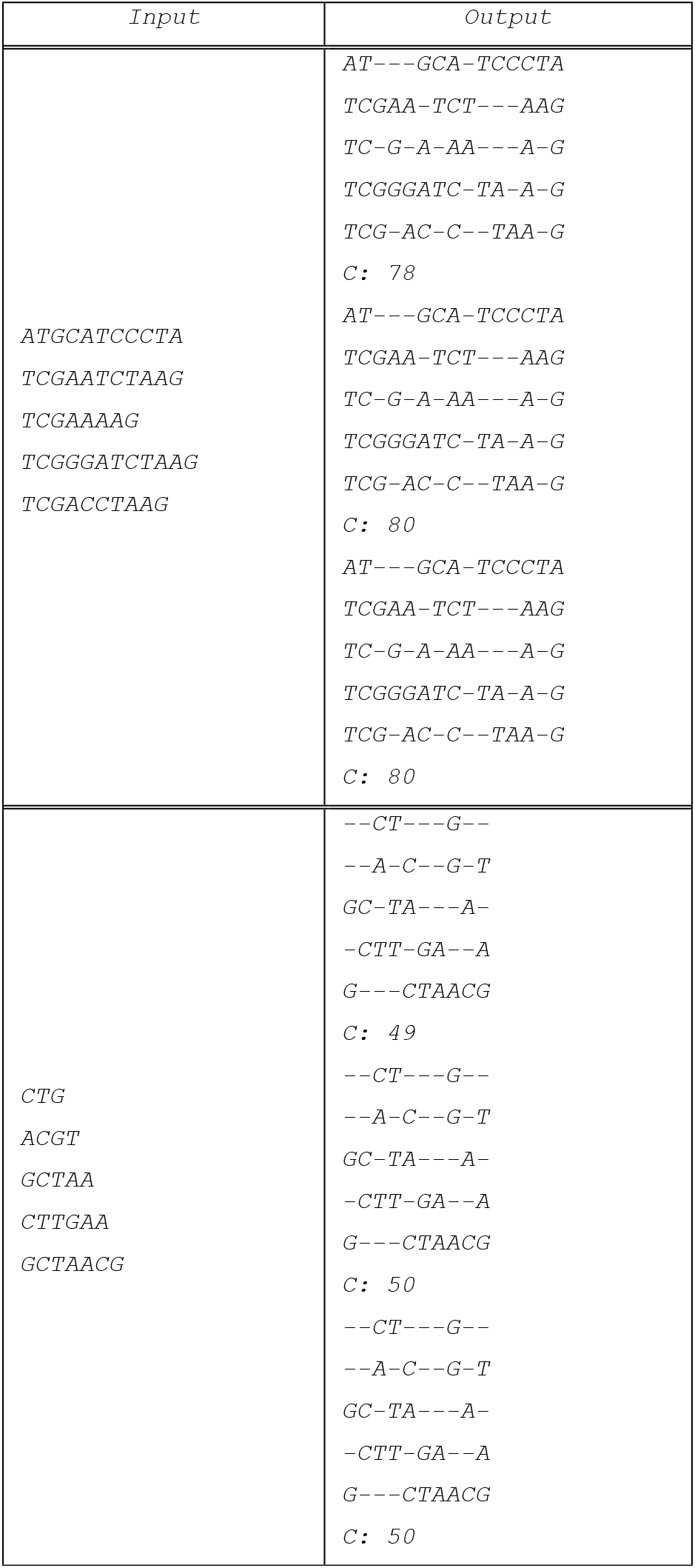
Some results from presented model

## 5. Discussion and Conclusion

The results indicate that the model succeeded in obtaining alignments in sequences of varying lengths that. Different results could be achieved by determining the gap penalty and substitution penalty and the appropriate function, depending on the researcher’s need.

Given that the matrix X covers all possible possibilities, the total space estimation will be 2^nm′^, in which the possible space will be equal to relation (8) as the relation (3) has limitations.

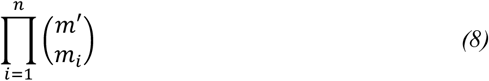

And, in case all the m_i_ values are the same, the relation will be as follows (9):

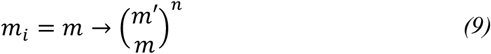

Moreover, the MSAs is mostly used in the approximately identical but long sequences, in which case a good result could be achieved by estimating the m′. In this case, if the input sequences are of the same length and the p-value is determined as small number, the Relation 10 will be established due to the high similarity of the sequences.

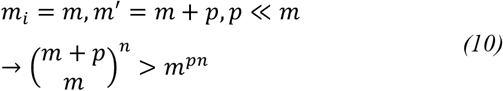

As explained in this relation, the number of possible states are definitely bigger than m^pn^ which mathematically proves that the MSA problem is NP-Complete.

Besides, based on the relation (6), if the string lengths are equal, we can obtain the overall optimal solution via relation (11):

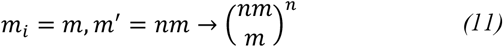

Based on this relation, the range for global optimal solution is very wide; this shows the researchers’ focus on heuristic approaches.

## Acknowledgement

I would like to express my special thanks and gratitude to my professor Dr. Kargari as well as Dr. Eshghi to helped me through in carrying out this study.

